# Alpha synuclein aggresomes inhibit ciliogenesis and multiple functions of the centrosome

**DOI:** 10.1101/2019.12.20.884643

**Authors:** Anila Iqbal, Marta Baldrighi, Jennifer N. Murdoch, Angeleen Fleming, Christopher J. Wilkinson

## Abstract

Protein aggregates are the pathogenic hallmarks of many different neurodegenerative diseases and include the Lewy bodies found in Parkinson’s disease. Aggresomes are closely-related cellular accumulations of misfolded proteins. They develop in a juxtanuclear position, adjacent to the centrosome, the microtubule organizing centre of the cell, and share some protein components. Despite the long-standing observation that aggresomes/Lewy bodies and the centrosome sit side-by-side in the cell, no studies have been done to see whether these protein accumulations impede the organelle function. We investigated whether the formation of aggresomes affected key centrosome functions: its ability to organize the microtubule network and to promote cilia formation. We find that when aggresomes are present, neuronal cells are unable to organise their microtubule network. New microtubules are not nucleated and extended, and the cells fail to respond to polarity cues. Since dopaminergic neurons are polarised, ensuring correct localisation of organelles and the effective intracellular transport of neurotransmitter vesicles, loss of centrosome activity could contribute to loss of dopaminergic function and neuronal cell death in Parkinson’s disease. In addition, we provide evidence that many cell types, including dopaminergic neurons, cannot form cilia when aggresomes are present, which would affect their ability to receive extracellular signals.

## Introduction

Parkinson’s disease is a progressive neurodegenerative condition that affects 1 in 500 of the population (Alves et al, 2008; Schrag et al, 2000; Van Den Eeden et al, 2003; von Campenhausen et al, 2005). Dopaminergic neurons in the substantia nigra pars compacta are first affected and the characteristic early symptom of Parkinson’s disease is tremor (Alves et al, 2008). As the disease progresses, other parts of the brain and nervous system are affected, with dementia occurring in later stages. Within neurons of Parkinson’s disease patients, large alpha-synuclein-positive intracellular inclusions known as Lewy bodies are observed (Wakabayashi et al, 2012). Increased α-synuclein (α-syn) levels, which occur in rare cases with multiplication of the SNCA gene encoding it, are sufficient to cause Parkinson’s disease (Ibanez et al, 2004; Singleton et al, 2003) and mutations associated with SNCA lead to an increase in aggregation propensity (Conway et al, 1998; Kruger et al, 1998; Polymeropoulos et al, 1997). These mutations cause α–syn to form oligomers and fibrils then aggregates. Large aggregates of α-syn constitute the Lewy bodies frequently found in the neurons of Parkinson’s disease patients (Baba et al, 1998; Spillantini et al, 1997). It is unclear if these aggregates or Lewy bodies are a means to protect the cell from smaller unfolded units of α–syn or if these structures cause neuronal death by obstructing the normal function of the cell.

Lewy bodies are observed in several diseases (Parkinson’s disease, dementia with Lewy bodies, incidental Lewy Body disease (Wakabayashi et al, 2012)) but are not found in healthy cells and are related to the aggresome, a structure found in cells that are processing large amounts of waste, unfolded polypeptide (Johnston et al, 1998). The aggresome is juxtanuclear inclusion body containing heat-shock proteins and components of the ubiquitin-proteasome system (Olzmann et al, 2008). These components are also found within Lewy bodies and there are shared ultrastructural similarities. This has led to the proposal that Lewy bodies are derived from aggresomes to specifically deal with misfolded α-syn (McNaught et al, 2002; Olanow et al, 2004).

The juxtanuclear location of the aggresome is shared by the centrosome, the microtubule organising centre of the cell. Indeed, they sit side-by side and centrosomal markers such as gamma tubulin are often used to detect the aggresome, alongside the intermediate filaments such as vimentin that cage the structure (McNaught et al, 2002). While the position of the aggresome at the centre of the microtubule network has logic in terms of transporting unfolded protein to a central location to be further processed (e.g. by proteasomal degradation), it may affect the function of the centrosome, whose major role is to organise this part of the cytoskeleton. Steric hindrance on a macromolecular scale may perturb centrosome function. Furthermore, the centrosome is used to make the cilium. This hair-like structure on the surface of cells has important roles both in motility, inter-cellular communication and monitoring the external environment, with very specialised cilia housing photopigments and olfactory receptors (Ibanez-Tallon et al, 2003; Nigg & Raff, 2009). To make the cilium, the centrioles of the centrosome migrate to the cell surface, a process that could be blocked by a large aggregate of protein smothering the centrosome or sticking it to the nucleus.

The colocalization of the aggresome and centrosome and the sharing of protein components has been known for nearly twenty years. However, it has not yet been tested if this colocalization affects the function of the centrosome. This could have important implications for the aetiology of Parkinson’s disease and other neurodegenerative diseases in which such aggregates are formed. In this study we sought to test whether multiple functions of the centrosome were impeded by the presence of aggresomes in their close vicinity using both *in vitro* and *in vivo* models. Our results suggest that inhibition of centrosome function might contribute to loss of function in neurons where there is aggregation of alpha synuclein.

## Results

### Aggresomes localise in close proximity to the centrosome

We confirmed that we could induce the formation of aggresomes in a variety of cell types using two previously published methods (Tanaka et al, 2004; Winslow et al, 2010). Cells were either treated with MG-132, a proteasome inhibitor, or transfected with expression constructs encoding GFP fusions of human α–syn wildtype or mutant versions, A30P and A53T, found in familial cases of Parkinson’s disease (Kruger et al, 1998; Polymeropoulos et al, 1997). The presence of aggresomes was then confirmed by staining with established markers for aggresomes: anti-vimentin or anti-gamma tubulin antibodies (Johnston et al, 1998; Olanow et al, 2004; Wigley et al, 1999). We tested aggresome formation in SH-SY5Y cells, a neuroblastoma line, either growing in continuous culture or after differentiation into dopaminergic neurons if treated with retinoic acid, and in rat basal ganglion neurons. Using both approaches, aggresome formation was observed, with vimentin caging the aggresomes and gamma tubulin seen as a dense area of staining next to the nucleus instead of the usual two punctae, representing the centrosome (Supplementary Fig. S1). Importantly, endogenous α-syn was observed in aggresomes induced by MG-132, demonstrating that both methods resulted in the accumulation of disease-associated, aggregate-prone proteins. These two methods were used in parallel in the majority of studies, however, low transfection efficiency prevented the use of the α–syn overexpression constructs in differentiated SH-SY5Y cells or rat basal ganglion neurons.

### Aggresomes suppress microtubule nucleation

The major function of the centrosome in interphase cells is the nucleation and organisation of the microtubule network (Bornens, 2002). The ability of the centrosome to nucleate microtubules can be assayed by the microtubule regrowth assay, in which the microtubules are first depolymerised by cold treatment for 30 min (Fig 1, 37°C, t-30 column vs 4°C, t = 0 column) followed by warming of the cells so that the centrosome can nucleate a new network (Fig 1, columns 37°C, t+0.5’ to 37°C, t+10’) (De Brabander et al, 1986; Fry et al, 1998). In control, untreated SH-SY5Y cells, the microtubules were nucleated after 30 s of warming following depolymerisation, with a clear aster of alpha tubulin staining and an extensive network after 10 min, as visualised by alpha tubulin staining (Fig. 1A, top row). In contrast, cells treated with 1µM MG-132 for 24 h prior to the assay did not nucleate any microtubules after 10 min warming (Fig 1A, second row), with only 5.3 ± 0.58% of treated cells starting nucleation but 84 ± 5.0% of untreated cells making an aster (Fig. 1B 100 cells, triplicate experiment). For SH-SY5Y cells transfected with expression constructs for GFP-tagged α-syn, only 23 ± 3.5% of the cells initiated microtubule nucleation by 10 min whereas 72 ± 3.8% of cells transfected with a control GFP expression plasmid re-established their network within the same time (Fig. 1A third row, transfected cells delineated with dashed line, identified by GFP expression (not shown); (GFP-transfected, familial mutant GFP fusions and same data with GFP signal from transfected cells all shown in Supplementary Fig. S2; 100 cells, triplicate experiment).

**Figure 1.**
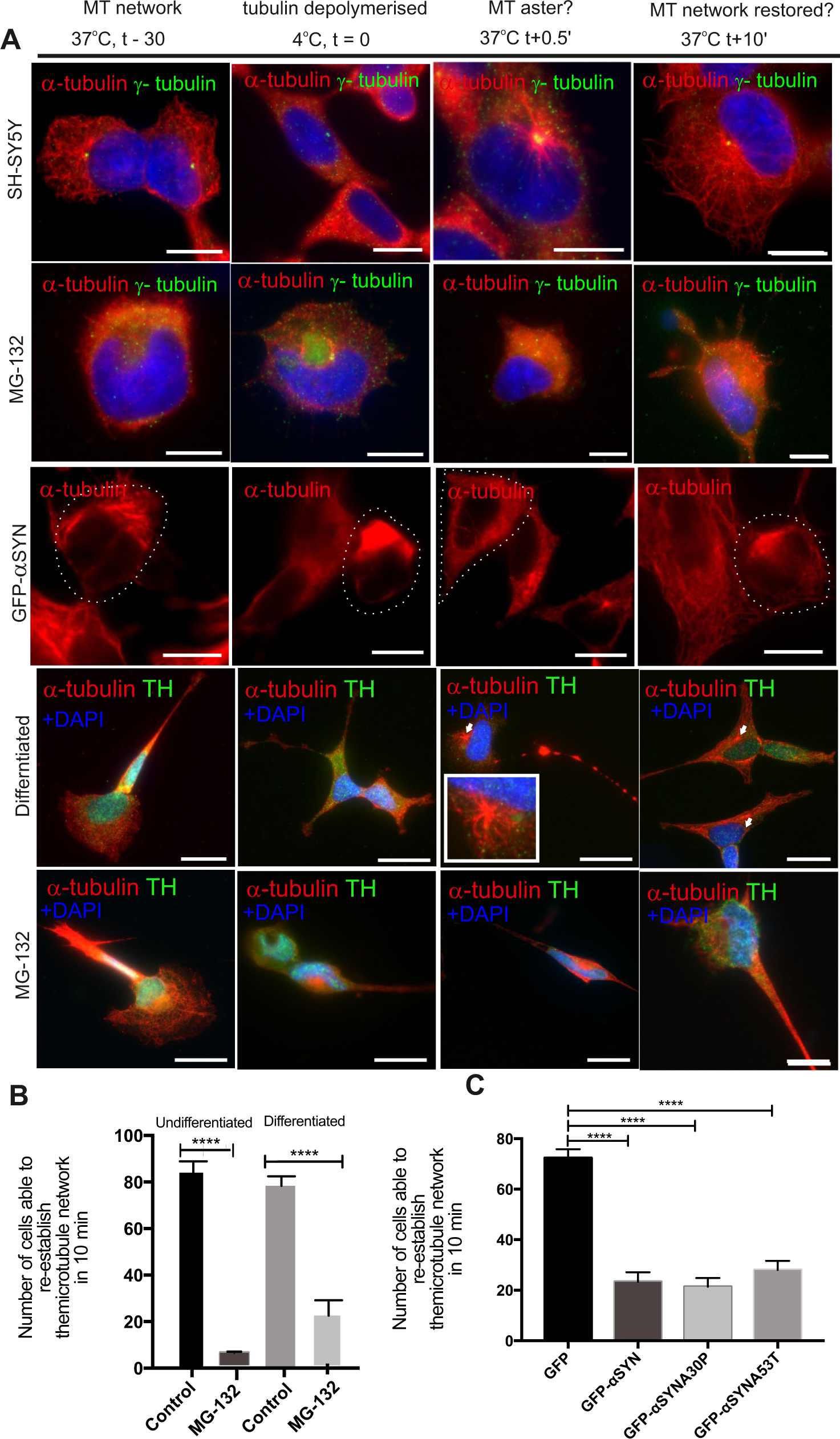
Microtubule nucleation is disrupted in the presence of aggresomes in undifferentiated and differentiated SH-SY5Y cells. A, top row) SH-SY5Y cells have an extensive microtubule network. Upon cold treatment microtubules depolymerise. Upon warming, microtubules nucleate from the centrosome forming a characteristic aster, which continues to grow until the network is re-established. In SH-SY5Y cells the aster is seen within in 30 seconds and the microtubule network is re-established within 10 min. A, row 2) In the presence of aggresomes (1μM MG-132 for 18 hours) the centrosome is unable to nucleate microtubules to re-establish this network. A, third row) Aggresomes formed by the overexpression of α-syn (GFP-fusion) had a similar affect: the centrosome is unable to re-establish the network in 10 min. A, row 4 and 5) In differentiated SH-SY5Y (tyrosine hydroxylase in green). microtubule nucleation is seen as asters form from the centrosome (arrow heads and inset). In the presence of aggresomes (1μM MG-132 for 18 hours) the density of the network is reduced and microtubule nucleation is severely compromised with staining of microtubules seen only in the last time point with no sign of asters forming. B) Quantification of microtubule regrowth in undifferentiated and differentiated SH-SY5Y when treated with MG-132 (p=0.0001, by Student’s t-test, 100 cells, n=3). C) Quantification of microtubule regrowth in SH-SY5Y when α-synuclein is over-expressed (p=0.0001, one way-ANOVA, 100 cells, n=3). Microtubule nucleation and re-establishment of this network was quantified by scoring cells (yes or no) whether the network was re-established in 10 mins.

Due to the morphology of the cell and the number of stabilized microtubules, the network and reforming aster is more difficult to observe in differentiated SH-SY5Y cells than in continuously growing cell lines. Nevertheless, whereas untreated cells were able to reform their labile microtubule network, differentiated SH-SY5Y cells were not, when aggresome formation was induced by MG-132 treatment (Fig. 1 bottom two rows). 21 ± 6.7% of treated cells were able to make an aster whereas 78 ± 4.2% of untreated cells re-establish their labile microtubule network. Quantification is show in Fig. 1B & C.

### Aggresomes prevent cell migration

The microtubule network is remodeled in response to polarity cues. This is exemplified by cell migration, during which the Golgi and centrosome are re-orientated to face the direction of cell locomotion. This function of the centrosome can be tested by the wound assay developed by Hall (Nobes & Hall, 1999). A strip of cells is removed from a confluent culture of an amenable cell line and those at the border of this wound will migrate to close the gap. We tested the ability of cells to migrate in the presence of aggresomes induced by MG-132. RPE1-hTERT cells generate aggresomes in the presence of 1 µM MG-132 as assayed by changes in gamma tubulin staining, from two punctae to a large area of signal close to the nucleus (Fig. 2: A, untreated; B, treated). RPE1-hTERT cells can migrate and close a ‘wound’ in 12 h (Fig. 2D-G). In the presence of aggresomes generated by MG-132 treatment, migration was halted with no cell movement to close the gap (Fig. 2H-K). Time-lapse observations of treated and untreated cells (Supplementary Videos 1 and 2) clearly demonstrate this lack of migration, quantified in Fig. 2C. The Golgi also did not re-orientate as in control cells, with control cells’ Golgi moving to face the wound (Fig. 2L vs M) whereas in treated cells the Golgi remained randomly orientated (Fig. 2N vs O). 25% of the wound was closed in treated cells versus near-complete wound closure for untreated cells. If we measure the angle of the Golgi relative to a line perpendicular the wound, then treated cells have a random orientation of the Golgi (average of 66.7° displacement from perpendicular) whereas control cells are facing the wound (average of 44.2° displacement from perpendicular) (Fig. 2P,Q).

**Figure 2.**
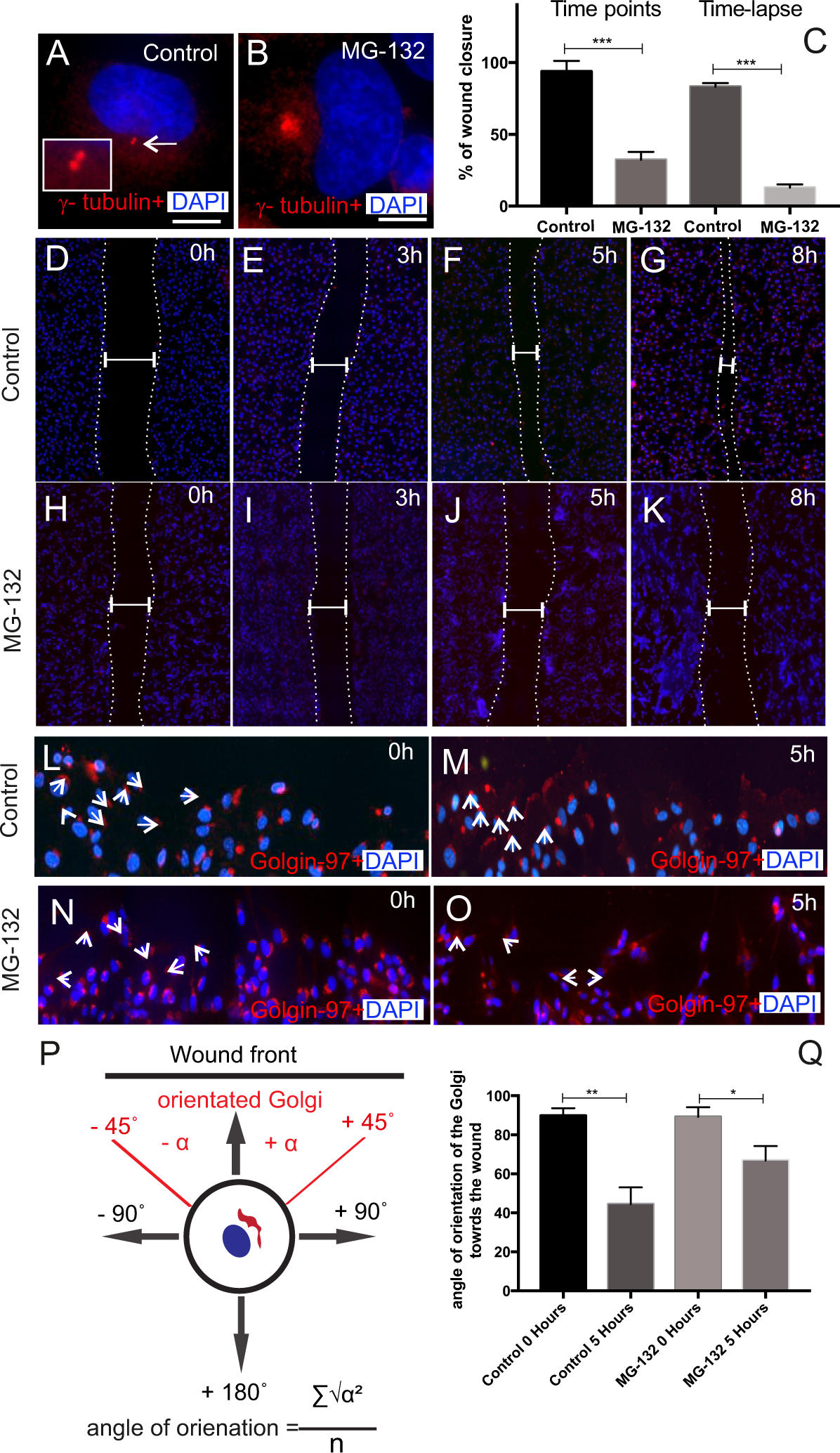
Aggresomes reduce rate of cell migration and inhibit polarity changes. A-B) In untreated RPE1-hTERT cells, γ-tubulin stains the centrosome. Upon MG-132 treatment (1μM for 18 hours) it stains the aggresome. C) The presence of aggresomes reduces the rate of cell migration in the scratch-wound assay. D-G) RPE1 cells are able to migrate when a strip of cells is removed from a monolayer of confluent cells. Complete wound closure is observed by 8 hours. H-K) In the presence of aggresomes, minimal cell migration is detected. L, M) In control cells the Golgi orientates from a random direction to face the leading edge of the wound (Golgin-97, red). N, O) In the presence of aggresomes, this change in orientation was not seen. P, Q) Quantification of change in angle of orientation during cell migration with a schematic diagram showing how the Golgi orientation was measured. (p=0.0011, by Student’s t-test, 100 cells, n=3). Scale bars 10μm. Nuclei stained with DAPI (blue).

### Aggresome formation inhibits ciliogenesis

We next tested if the presence of aggresomes prevents cells from making cilia as ciliogenesis requires the centrioles of the centrosome to move to the cell surface where one centriole templates the axoneme that forms the internal structure of the cilium. Many cell types ciliate, but not all. To test whether aggresome inhibition of ciliogenesis might be relevant to the dopaminergic neurons affected early in Parkinson’s disease, we tested several cell types for the presence of cilia by staining for acetylated tubulin together with gamma tubulin to mark the basal bodies. Undifferentiated SH-SY5Y cells generated cilia when serum starved and GFP expression did not affect ciliogenesis (Fig. 3A,B). Undifferentiated SH-SY5Y cells failed to ciliate when treated with MG-132 (Fig. 3C) or when GFP-α-syn (any variant, only wild-type α-syn shown) was overexpressed (Fig. 3D). Quantification is shown in Fig 3I.

**Figure 3.**
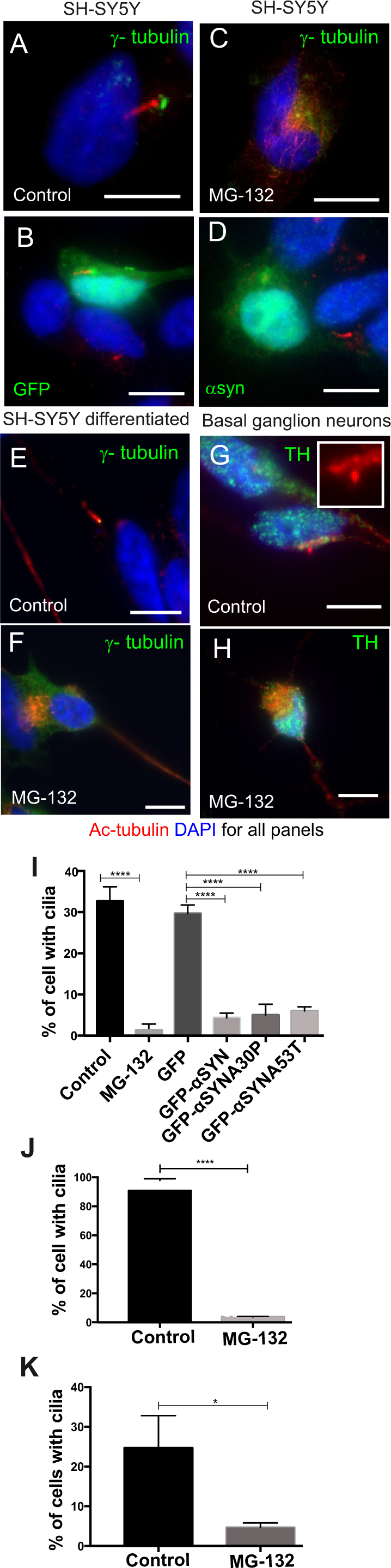
Aggresomes inhibits ciliogenesis. A, B) Undifferentiated SH-SH5Y cells form cilia in serum free conditions or when transfected with a control GFP only expression plasmid. C, D) When treated with MG-132 (1μM for 18 hours) or transfected with a GFP-α-syn expression plasmid, cilia formation is inhibited. E) Differentiated SH-SY5Y cells form cilia in serum free conditions. F) In the presence of aggresomes (MG-132 1μm for 18 hours) cilia are unable to form. γ-tubulin stains the aggresomes. G) Cilia are also found in TH-positive basal ganglion neurons, acetylated tubulin in red, TH in green. H) When treated with MG-132 (3μM for 18 hours) cilia are no longer visible. I) Quantification of ciliation in undifferentiated SH-SY5Y cells under various treatments (untreated vs. MG-132, p=0.0001, by Student’s t-test, 100 cell, n=3; GFP expressing vs. α-syn expression, p=0.0003, by one-way ANOVA, 100 cells, n=3). J) Quantification of cilia in differentiated SH-SY5Y cells when treated with MG-132 (p=0.0001 by Student’s t-test, 100 cells counted, n=3. K) Quantification of cilia for basal ganglion neurons when treated with MG-132 (p=0.0126 by Student’s t-test, 100 cells counted, n=3). Scale bars 10μm. DNA/nuclei stained with DAPI (blue).

Differentiated SH-SY5Y cells formed cilia as part of their differentiation programme (Fig. 3E) but did not do so when treated with MG-132 (Fig. 3F). Rat basal ganglion cells are ciliated under their normal culture conditions (Fig. 3G) but their ciliogenesis was much reduced when treated with MG-132 (Fig. 3H). Quantification is shown in Fig. 3J,K.

To test whether aggresome formation affected ciliogenesis in an *in vivo* system, we performed α-syn over-expression experiments in zebrafish embryos. Cilia are abundant in zebrafish embryos and larvae and the olfactory pit in embryonic / larval zebrafish is highly ciliated and accessible to observation. The ciliated cells of the olfactory pit are dopaminergic neurons, as indicated by tyrosine hydroxylase (TH)-positive staining (Fig. 4 A). Treatment of embryos with MG-132 caused loss of cilia at 3 dpf (Fig. 4: B, untreated; C, treated) with cilia number being reduced from 65 ± 11 per pit to 15 ± 6 (Fig. 4I). We injected 1-cell embryos with *in vitro* transcribed mRNA encoding human α–syn, (WT and familial mutants) and GFP as a control (Fig. 4 D, E). Larvae were fixed at 2 dpf and assayed for olfactory cilia by wholemount acetylated tubulin staining (Fig. 4 F, G; again familial mutants show same result as WT α–syn). Cilia numbers were much reduced (11 ± 3.6 vs 41 ± 5.3 per pit) in the olfactory zone when the embryos overexpressed α–syn, any variant (Fig. 4H). The remaining cilia were reduced in length, by approximately 50% from 8.4 ± 0.49 µm to 4.7 ± 0.71 µm (Fig. 4J). The gross anatomy of these embryos was otherwise normal (Fig 4D, E; data not shown). Absence of cilia may be expected to cause other defects during embryogenesis, for example hydrocephaly, left-right asymmetry defects and pronephric cysts. We did not observe these defects. However, the effect on ciliogenesis at the olfactory pits was mild at 24 hpf. We therefore suspect that accumulation of α-syn is required over several days to inhibit ciliogenesis and so earlier stages escape the effects of aggresome-induced cilium loss.

**Figure 4.**
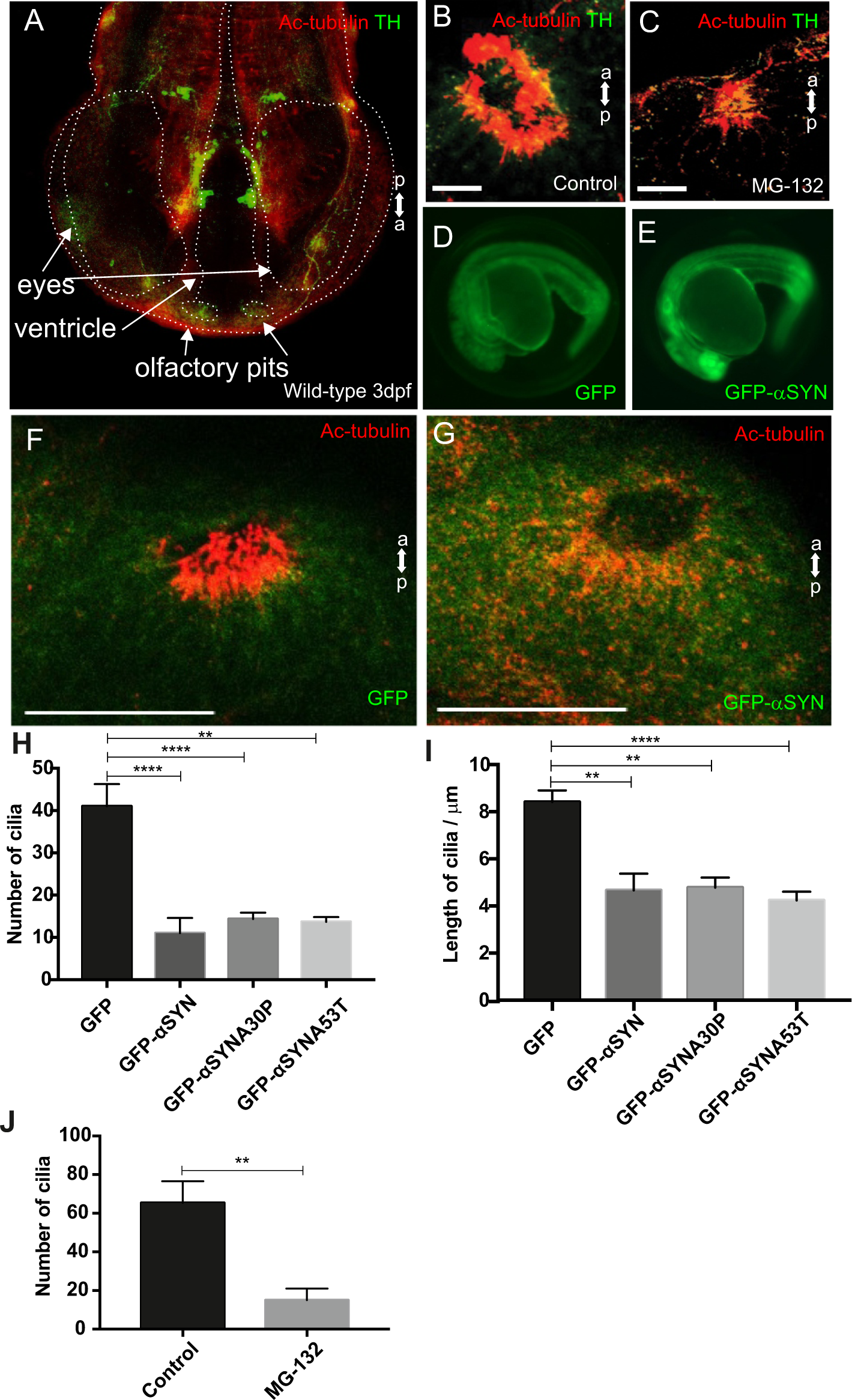
Olfactory cilia in zebrafish embryos are severely reduced in the presence of aggresomes. A) The neuronal dopaminergic network in 3 d.p.f. zebrafish forebrain viewed from the dorsal aspect, detected by TH-staining (green). TH staining is also seen around the olfactory pit. Acetylated tubulin (red) stains axon tracts and cilia. B) By 3 d.p.f extensive numbers of cilia are visible at the olfactory pit. C) Embryos treated with MG-132 (50μM for 48 hours), showed an extensive reduction in number of cilia. D, E) Overexpression of control GFP or α-synuclein does not cause any anatomical defects in zebrafish larvae. F) In control GFP-injected embryos, cilia are seen in the olfactory pit in large numbers. G) Overexpression of any of the three forms of α-syn severely reduces numbers of cilia. Cilia length is also reduced. H) Quantification of cilia numbers in zebrafish embryos when α-syn is over expressed (p=0.001, by one-way ANOVA, n=3,). I) Quantification of length of cilia when α-syn is overexpressed in zebrafish embryos (p=0.0088, by one-way ANOVA, n=3). J) Quantification of cilia numbers when embryos are treated with MG-132 (p=0.0024, by Student’s t-test, n=4. Scale bar 100μm.

## Discussion

Aggresomes occupy a central position in the cell, at the hub of the microtubule network and that places them in the vicinity of the centrosome, the major microtubule organising centre of the cell. Furthermore, the aggresome acquires an important centrosomal component, gamma tubulin. We show here that aggresomes severely compromise centrosome function. Microtubule nucleation is severely reduced and the centrosome is unable to be repositioned during cell migration. These affects are observed whether the aggresomes are generated by proteasome inhibition or alpha synuclein overexpression. Indeed MG-132 treatment generates aggresomes in which endogenous alpha synuclein accumulates. These results and are consistent with and extend previous observations of the effect of MG-132 on microtubule nucleation (Didier et al, 2008). A simple, steric hindrance of the centrosome is the simplest and most likely explanation for this effect.

The centrosome is the main microtubule organising centre in the cell, nucleating microtubules and anchoring a portion near to the nucleus. Its role as a major component of the spindle poles is not important in mature neurons which do not divide. However, a functioning, polarised microtubule network is still required for intracellular trafficking in a terminally differentiated cell such as neurons and the centrosome is essential for maintaining this. Much research on Parkinson’s disease currently focusses on the mitochondrion (Exner et al, 2012). Without a functioning centrosome and therefore a properly organised microtubule network, it is likely that the mitochondria will themselves not be trafficked correctly in the cell, with a particular impact on cells with a high energy demand, like neurons.

Most neurons do not migrate so the effect of aggresomes on cell migration is probably not relevant to the development of Parkinson’s disease. However, the inability of cells to migrate and the Golgi to re-orientate in this situation provides a useful surrogate read-out for the inability to repolarise the microtubule network in a desired direction. In mature neurons, compromised cell polarity could have severe long-term effects on neuronal function and survival. A polarised microtubule network is essential for the rapid trafficking of organelles and vesicles, especially those containing neurotransmitters that need to be transported to synapses, and the recycling of materials involved in neurotransmission.

Nearly all cell types make a cilium but neurons are one group of cells that contain both unciliated (and centrosome possessing) sub-types and ciliated (acentrosomal) subtypes. Our results show that aggresomes can prevent a cell from turning its centrosome into a cilium. In Parkinson’s disease, this may be relevant to those cells that undergo ciliogenesis. An early symptom of Parkinson’s disease is loss of smell and anosmia may precede other symptoms (Doty et al, 1988; Haehner et al, 2007). It is not clear what the cause of anosmia is, the assumption being that loss or impairment of neurons processing olfactory information is the cause. An alternative explanation is that the olfactory neurons themselves are affected during Parkinson’s disease progression. Olfactory neurons are part of the dopaminergic system (Pignatelli & Belluzzi, 2017) and the olfactory receptors are housed in cilia on these cells. If early in the development of Parkinson’s disease the ability of olfactory neurons to renew cilia was compromised then so would the sense of smell. It may be possible, then, that this early symptom is a result of gradual loss of cilia from olfactory neurons. If this is the case, then cilia density in the olfactory epithelium of Parkinson’s patients should be reduced in age-matched controls. This hypothesis certainly warrants further investigation as it may reveal that the olfactory cells are the sentinels of the dopaminergic system and anosmia represents a first indicator of dopaminergic cell dysfunction. Should olfactory cilia be accessible to routine screening, their number may be an early diagnostic tool for Parkinson’s disease.

While we have provided clear evidence that aggresome formation affects centrosome function in cell and *in vivo* models, proving that Lewy bodies affect microtubule nucleation and cellular polarity in neurons of Parkinson’s patients is likely to prove difficult. However, it may be feasible to assay some of these effects in patient fibroblasts, which are more readily accessible. Furthermore, since microtubule regrowth is a biochemical phenomenon whose components may be capable of withstanding freezing it may be technically possible to observe this in clinical samples obtained from brain bank tissue.

## Materials and Methods

### Cell Culture

Cell lines used include: HeLa, provided by Prof George Dickson at Royal Holloway; neuroblastoma cell line SH-SY5Y, provided by Prof Robin William at Royal Holloway; immortalised human retinal pigment epithelial cells (RPE1-hTERT), kindly provided by Prof. Erich Nigg, Basel, Switzerland; mouse embryonic fibroblast cells (MEFs), provided by Dr Jenny Murdoch at Royal Holloway. Primary rat basal ganglion neurons were prepared by Dr Simona Ursu at Royal Holloway.

HeLa and SH-SY5Y cells were maintained in Dulbecco’s Modified Eagle’s Medium (DMEM) supplemented with 10% Foetal Bovine Serum (FBS), 1x Antibiotic-Antimycotic mix (Gibco; 5000 units of penicillin, 5000μg streptomycin and 15μg Amphotericin B) and 2mM L-glutamine. RPE1-hTERT were grown in Dulbecco’s Modified Eagle’s Medium with nutrient mixture F-12 Ham (DMEMF-12) supplemented with 10% FBS, 1x Antibiotic-Antimycotic (5000 units of penicillin, 5000μg streptomycin and 15μg Gibco Amphotericin B), 2mM L-glutamine and 0.38% Sodium bicarbonate. MEFs were grown in DMEM supplemented with 10% FBS, 1x Antibiotic-antimycotic, 2mM L-glutamine and 1x non-essential amino acids. All cell lines were cultured at 37°C in a humidified atmosphere with 5% CO_2_. The same media mix was used for future experimental assays unless otherwise stated.

### Primary Basal Ganglion neurons

Primary cultures of basal ganglion neurons were prepared from E18 Sprague Dawley rat embryos as previously described (Marsh et al, 2017). Briefly, cells were plated at a density of either 75,000 or 500,000 cells on poly-D-lysine-coated (Sigma, 0.1 mg/ml in borate buffer pH 8.5) 22mm^2^ glass cover slips. The plating medium was DMEM supplemented with 5% FBS, penicillin/streptomycin and 0.5 mM L-glutamine (all from Invitrogen). On the next day the medium was changed to full Neurobasal medium (Neurobasal medium supplemented with B27, 0.5 mM L-glutamine, all from Invitrogen). Cultures were incubated at 37 °C and 5% CO_2_, and were used at 18 days *in vitro*.

### SH-SY5Y Differentiation

Cells were plated on collagen-coated coverslips in 12-well plates. Optimised seeding density was calculated to be 6 × 10^4^ cells in 12-well culture plates. Cells were seeded out in DMEM serum supplemented media, The media was replaced the following day with DMEM-F12 supplemented with 1% FBS, 2mM L-Glutamine, 1x Antibiotic-Antimitotic mix (5000 units of penicillin, 5000μg streptomycin and 15μg Gibco Amphotericin B), 1 X non–essential amino acids and 10 μM all trans-retinoic acid (RA). Every two days the media was replenished and after 7 days differentiation was observed.

### Aggresome formation

Aggresomes were induced by either treating cells with the proteasome inhibitor MG-132 (Sigma Aldrich) or by overexpressing GFP-tagged human α-syn. The optimal MG-132 concentration was determined for each cell line ranging from 1μM to 10μM (HeLa, 10 μM; SH-SY5Y, 1 μM; differentiated SH-SY5Y, 1μM; MEFs, 5 μM and rat neurons, 3 μM). MG-132 was added to media for 18 hours at 37°C in a humidified atmosphere with 5% CO_2_.

Constructs including peGFP-C2, peGFP-αSYN, peGFP-αSYNA30P and peGFP-αSYNA53T were transiently transfected into cells. Lipofectamine 2000 (ThermoFisher) was used as the transfection reagent following manufacturer’s instruction. In brief, cells were plated onto glass coverslips in serum-supplemented media without antibiotic-antimycotic mix. At 70% confluency cells were transfected with DNA-lipid complexes (2.5 μg of plasmid DNA was diluted in 250 µL Opti-MEM + 10 μL of Lipofectamine 2000 reagent in 240 μL of Opti-MEM). Complexes were added to each well for 5 hours at 37°C after which the media was replaced by low serum-supplemented media (DMEM, 1 % FBS, 2mM L-glutamine and 1% antibiotic-antimitotic mix for HeLa and SH-SY5Y; DMEM-F12, 1 % FBS, 1% antibiotic-antimitotic mix for RPE1-hTERT). Overexpression was achieved at 72 hours from the time of transfection.

### Microtubule re-growth assay

Cells were plated onto ethanol-washed glass coverslips in serum-supplemented media. At 70% confluency the samples were processed for the respective condition (MG-132 treatment or overexpression of α-syn). The plate was incubated on ice for 30 min then pre-warmed media (37°C) was added. Samples were fixed at different time points ranging from 0.5 minutes to 25 minutes, depending on the cell line, including immediately after 30 min on ice. Cells were fixed for 10 min with methanol (−20°C), washed with PBS and stored in PBS.

### Ciliogenesis assay

Cells were plated on glass coverslips in 6- or 12-well culture plates in serum-supplemented media. Cells were treated with MG-132 or transfected with α-syn overexpression constructs to form aggresomes. The media was replaced the following day with serum free media; samples were incubated for 24 hours to induce cilia formation. Cells were fixed using 4% (v/v) formaldehyde (FA) for 10 min. FA was aspirated and cells were washed with PBS. Samples were stored in 0.2 % Triton in PBS until they were processed for immunocytochemistry.

### Cells migration assay (Scratch-Wound Assay)

MEFs and RPE1-hTERT were used in the wound assay. MEFs were plated onto collagen-coated 35mm plastic Petri dishes. At 100 % confluency (with or without aggresome formation using MG-132) a P200 pipetman tip was used to make wound. The media was aspirated and washed twice with 1x PBS to remove detached cells. CO_2_ independent media supplemented with 10% FBS, 2mM L-glutamine, 1 x antibiotic-antimycotic mix was added for time-lapse experiments. Images were taken using a Nikon TE300 microscope with a 37°C chamber, over a 24 hour period with images taken every 2 min. Similarly RPE1-hTERT cells were plated onto ethanol-washed glass coverslips and treated with MG-132. Cell migration was assessed by fixing cells at set time points with ice-cold methanol.

### Golgi orientation

A Golgi positioned within-45° and +45° of the wound was considered to be orientated towards the wound. The average angle of orientation was calculated using the formula 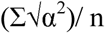 where α is the angle between the Golgi and a line perpendicular to the wound edge.

### Immunocytochemistry

Cells were fixed with either methanol (−20°C) or 4% (v/v) FA for 10 min. The fixative was removed and washed with PBS (3 × 5 min). Cells were blocked in 3% BSA for 30 min at room temperature. Cells were incubated with primary antibody in 1% BSA, either for 3 hours at room temperature on a shaking platform or overnight at 4°C on a shaking platform. After primary antibody incubation the cells were washed with PBS (3 × 5 min) and incubated with secondary antibody for 1 hour at room temperature on a shaking platform. Cells were washed with PBS (3 x 5 min), and mounted using FluorSave (Calbiochem). Primary antibodies used were: rabbit anti-Tyrosine Hydroxylase (Merck-Millipore), 0.1 μg/ml; mouse anti-vimentin (Sigma), 1μg/ml; mouse anti-acetylated-tubulin (Invitrogen) 1μg/ml; mouse anti-γ-tubulin (Sigma), 1μg/ml; rabbit anti-γ-tubulin (Sigma), 1μg/m; mouse anti-α-tubulin (Sigma), 1μg/ml; rabbit anti-α-synuclein (Cell Signalling), 10μg/ml; mouse anti-Golgin 97 (ThermoFisher), 1μg/ml. Secondary antibodies used were: goat anti-mouse IgG, Alexa Fluor 594-conjugated (Invitrogen) 1μg/ml; goat anti-mouse IgG, Alexa Fluor 488-conjugated (Invitrogen) 1μg/ml; and goat anti-rabbit IgG Alexa Fluor 488-conjugated (Invitrogen) 1μg/ml. Images were taken using a Nikon Ni-E fluorescence microscope.

### Whole mount immunostaining

Embryos were fixed with Dent’s fixative (80:20, Methanol: DMSO) or 4% (v/v) FA overnight at 4 °C. Fixative was removed the following day; embryos fixed with Dent’s fixative were stored in methanol at −20°C; FA fixed embryos were stored in PBS +0.2% Triton. Embryos fixed with Dent’s fixative were permeabilised by incubation in 100% methanol for 30 mins at −20°C. Embryos were rehydrated by washing in serial dilution of methanol in PBS including: MeOH:PBS at 70:30, 50:50 and 30:70 and final wash with PBS. FA fixed embryos were permeabilised by incubation in 0.25% trypsin-EDTA in PBS for 10 min on ice and then washed three times for 30 min in PBS +0.02% Triton. Embryos were blocked for 4 h in 10 % heat-inactivated goat serum, 1% bovine serum albumin and 0.2% Triton in PBS. Embryos were incubated with primary and secondary antibodies for 36 h in blocking solution. Primary antibodies used: rabbit anti-Tyrosine Hydroxylase (MERK Millipore), 0.1 μg/ml; and mouse anti-acetylated-tubulin (Invitrogen) 1μg/ml. Secondary antibodies used were goat anti-mouse IgG, Alexa Fluor 594-conjugated (Invitrogen) 1μg/ml; and goat anti-rabbit IgG Alexa Fluor 488-conjugated (Invitrogen) 1μg/ml. Confocal stacks were imaged with an Olympus FX81/FV1000 laser confocal system using Ar gas laser and He-Ne diode laser. Stacks were taken in 1μm thickness and are represented as maximum-intensity projections. Stacks were analysed using ImageJ.

### mRNA synthesis

mRNAs were transcribed from the Sp6 promoter of the pCS2+-based plasmids encoding α-syn and mutant forms, using the mMessage mMachine in vitro transcription kit (Ambion, TX, USA). RNA was purified using the Qiagen RNeasy kit (Qiagen).

### Zebrafish

Zebrafish were maintained and bred at 26.5°C; embryos were raised at 28.5°C. Both AB and TL wild-type strains were used for these studies. Embryos were processed by 3.d.p.f. No protected species, as defined by the Animals (Scientific Procedures) Act, 1986 were used for experiments in this study. Embryos were injected into the yolk with mRNA using a micromanipulator-mounted micropipette (Borosil 1.0 × 0.5 mm, Frederick Haer & Co., Inc., USA) and a Picospritzer microinjector. Between 150-200pg of mRNA was injected into the yolk of the embryos at 1-4 cell stage. For MG-132 treatment, embryos were treated with 50μM for 48 hours. Embryos processed for immunostaining were grown in 0.003% phenylthiourea to inhibit melanin production. mRNA synthesis described below.

### Statistics

Statistical tests used for each experiment are given in the figure legends. t-tests were used when comparing two treatments and ANOVA for multiple treatments. n numbers represent the experimental unit, either number of embryos or separate cell culture experiments.

### Animal studies

No protected stages of zebrafish, as defined by the Animals (Scientific Procedures) Act, 1986, were used for experiments in this study, only fry less than 5 d.p.f. Rats were killed by Schedule 1 methods, according to Home Office regulations, in compliance with the Animals (Scientific Procedures) Act, 1986.

## Author contributions

AI performed the experiments under the supervision of CJW and JNM. AI prepared the figures and legends. CJW wrote the main text. AF helped with study design and manuscript preparation.

## Acknowledgements

We thank Dr Simona Ursu for assistance with the primary neuronal cultures.

---

**Supplementary Figure S1.**
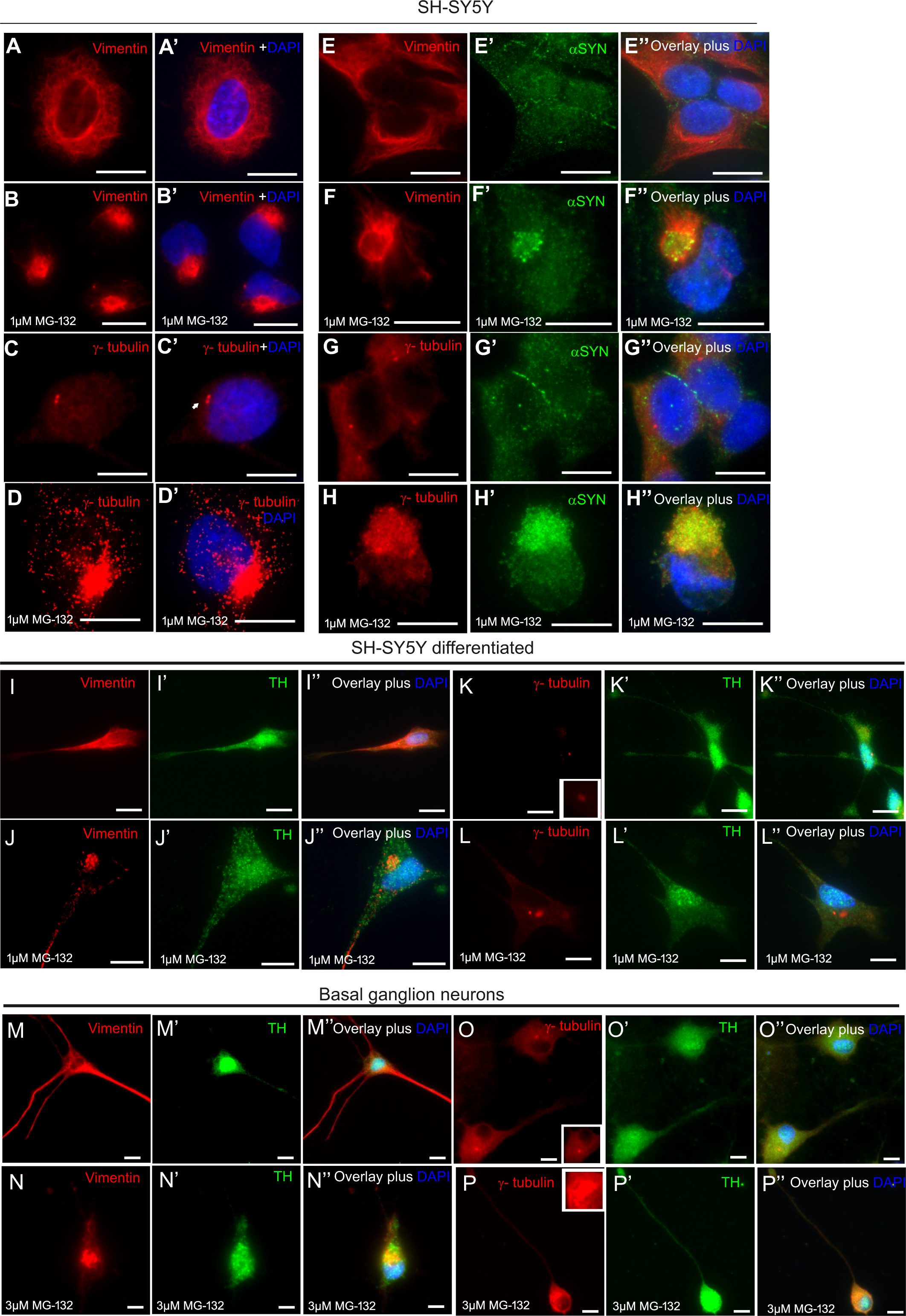
SH-SY5Y cells form aggresomes when treated with MG-132. Vimentin (A-A’) (red) in control HeLa cells forms a fibrous network, around the nuclei (DAPI-blue). (B-B’) When treated with 10μM MG-132 for 18 hours, vimentin positive aggresomes appear in cells, juxtaposed to the nucleus. The vimentin stain changes from a filamentous stain around the nuclei to caging the aggresome. Similarly, γ-tubulin (red) staining in control cells (C-C’) labels the centrosomes as two punctae. Following MG-132 treatment, γ-tubulin forms a condensed structure around the aggresome (D-D’). The expression pattern of endogenous α-syn was also investigated to determine whether this protein co-localises within the aggresome. (E’-E’’) In control cells, α-syn (green) staining was widespread and diffuse within the cytoplasm with vimentin (red) forming a filamentous network. (F’-F’’) Following MG-132 treatment, α-syn aggregates were observed and co-localised with vimentin staining within the aggresomes. (G’-G’’) In control cells, γ-tubulin (red) was observed as two punctae with α-syn diffuse within the cytoplasm. (H-H’’) γ-tubulin staining (red) also co-localises with endogenous α-syn in the aggresome when treated with MG-132. Differentiated SH-SY5Y cells were mock-treated (I-I”) and vimentin (red) staining was observed surrounding the nuclei as well as along the axon. J-J”) Cells treated with MG-132 (1μM for 18 hours) vimentin staining changed to a compact structure near the nuclei, indicative of aggresomes. K-K”) In mock-treated cells γ-tubulin formed two punctae next to the nucleus. L-L”) In treated cells, aggresomes were detected by γ-tubulin staining. Differentiated SH-SY5Y cells are TH positive. M-M’’) In rat basal ganglion neurons, vimentin staining (red) is abundant around the nuclei and along the axon. N-N’’) When treated with MG-132, vimentin localises to the aggresome. (O-O’’) In rat basal ganglion neurons, the γ-tubulin is observed at two punctae close to the nucleus. P-P’’) Upon MG-132 treatment, the γ-tubulin staining now forms a larger structure next to the nucleus.

**Supplementary Figure S2.**
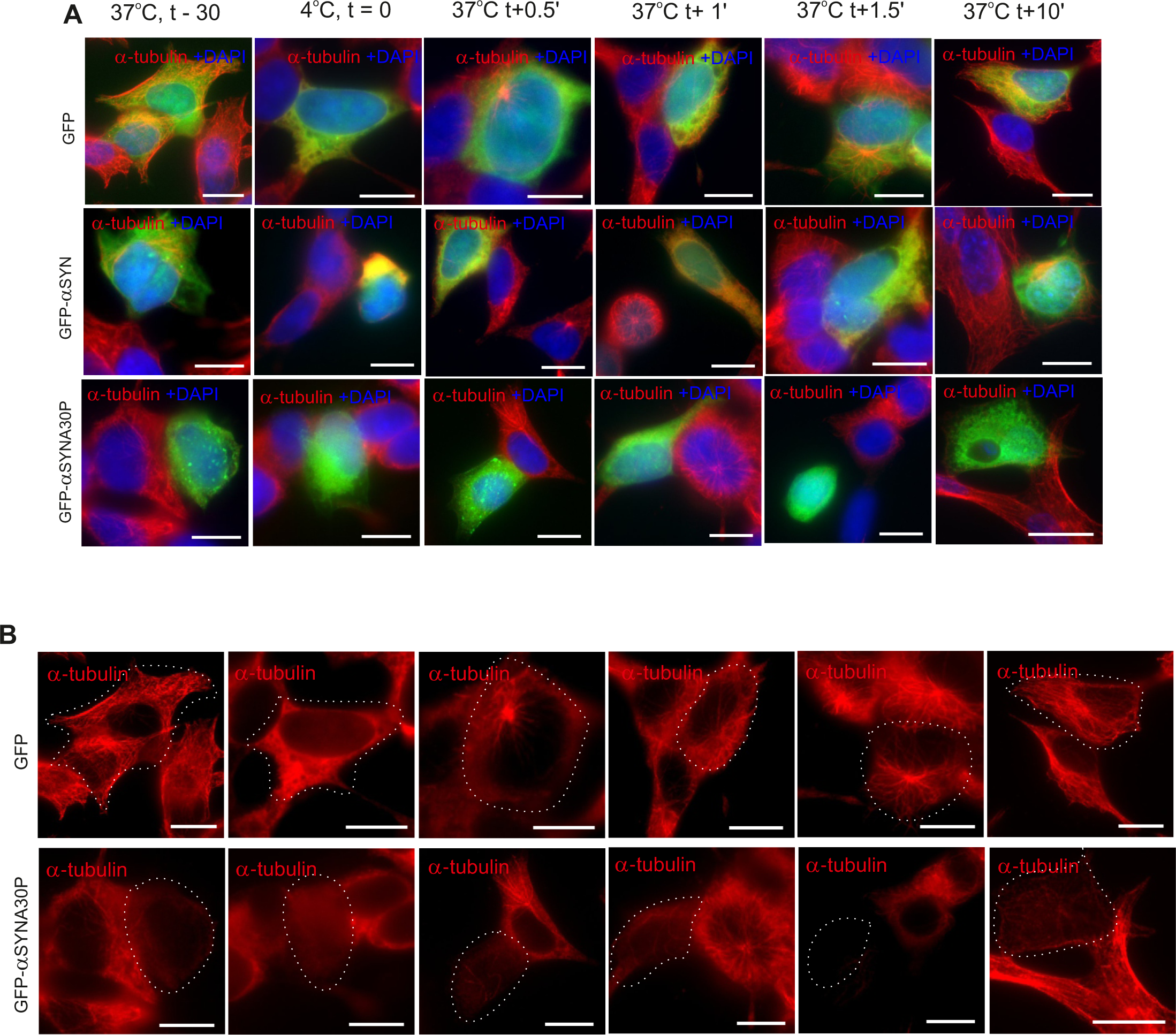
Microtubule nucleation is disrupted in the presence of aggresomes in undifferentiated SH-SY5Y cells. A) Top row, SH-SY5Y cells have an extensive microtubule network. Upon cold treatment microtubules depolymerise. Upon warming microtubules nucleate from the centrosome forming a characteristic aster, which continues to grow until the network is re-established. In SH-SY5Y cells transfected with GFP, the aster is seen within 30 seconds and the microtubule network is re-established within in 10. A) Second and third rows: aggresomes formed by the over expression of GFP-α-synuclein (α-SYN WT or αSYNA30P) inhibited microtubule nucleation: the centrosome is unable to re-establish the network in 10 min. Nuclei stained with DAPI (blue). B) Panels for first and third rows of A shown with alpha tubulin staining only. Scale bar 100μm.

**Supplementary Movie S1. Time-lapse of RPE1-hTERT migrating to close the wound**. Bright field view showing RPE1-hTERT cells migrate to close the wound when a strip of cells is removed from a monolayer of confluent cells within in 12 hours.

**Supplementary Movie S2. Time-lapse of RPE1-hTERT showing cells fail to close the wound in the presence of aggresomes.** Bright field view of RPE1-hTERT cells failing to migrate and close the wound in the presence of aggresomes induced by MG-132 exposure.

